# Cell barcoding with tandem fluorescent proteins enables high-throughput signaling dynamics analysis

**DOI:** 10.1101/2025.06.17.660155

**Authors:** Jhen-Wei Wu, Jr-Ming Yang, Suyang Wang, Yun Chen, Chuan-Hsiang Huang

## Abstract

Cell barcodes are essential for a wide array of experimental applications, including lineage tracing, genetic screening, and single-cell analysis. An optimal barcode library would provide high diversity, live-cell compatible identification, and simple readout. In this work, we introduce single chain tandem fluorescent protein (sctFP) barcodes, constructed by linking different fluorescent proteins (FPs) into a single polypeptide chain with varied copy numbers. We found that the fluorescence signal intensity ratio at different wavelengths can reliably differentiate sctFPs generated using cnidarian FPs, but not prokaryotic FPs that require exogenous cofactors. The sctFPs enable the multiplexing of genetically encoded fluorescent biosensors, enhancing current biosensor multiplexing methods through a simplified imaging and analysis pipeline that support high-throughput applications. Their robust spectral profiles are compatible with a broad range of biosensor types. Using sctFPs, we demonstrate simultaneous tracking of various signaling activities with biosensors of different spectral properties. Together, this strategy provides a robust and scalable method for barcoding cells across diverse experimental contexts.

## Background

Cell barcodes play a vital role in a variety of experimental applications focused on monitoring individual cells [1–4]. These unique identifiers allow researchers to map clonal relationships within cell populations and track individual cells over time. For example, barcodes enable the tracing of cell lineages and simultaneous evaluation of multiple genetic modifications to accelerate the discovery of functional gene targets [5–9]. For single-cell analysis, barcoding enables the differentiation and tracking of different cells, providing comprehensive insights into cellular heterogeneity and the dynamics of complex biological systems [10,11].

Cell barcoding is typically achieved using either nucleotide-based or fluorescent barcodes. Nucleotide-based barcodes take advantage of the large number of combinations that can be generated from relatively short nucleotide sequences [1,12,13]. However, nucleotide-based barcodes are in general destructive when identifying cells. Cell lysis and sequencing, amplification, or hybridization assays are required, thus are suitable for applications where the complexity and depth of information outweigh the need for dynamic and longitudinal observation. In contrast, fluorescent barcodes utilize the spectral and spatial patterns of fluorophores to identify different cell populations [14–18]. Barcodes based on fluorescent proteins (FPs) permit longitudinal monitoring of dynamic cell behaviors and phenotypes.

A recent application of FP-based barcodes is multiplex imaging of genetically encoded fluorescent biosensors [19–22]. In these applications, cells expressing different biosensors are tagged with barcodes created from combinations of FPs that are spectrally distinct from the biosensors. To mitigate the reduction of available spectral space due to biosensor fluorescence, barcoding FPs are directed to different subcellular locations. However, this localization-based barcoding approach presents several challenges. Firstly, identifying the barcodes often relies on high-resolution confocal imaging, making it difficult to apply to lower magnification images used in high-throughput screening. Additionally, distinguishing subcellular structures becomes challenging for cells with minimal cytoplasm (e.g., leukocytes) or those prone to clustering. These difficulties are often compounded by uneven expression levels of barcoding proteins, posing further obstacles to automating image analysis.

In this work, we explored a new strategy for generating a large number of FP-based barcodes with robust color patterns for labeling various cell populations, which can be combined effectively with fluorescent biosensors. This approach, termed single-chain tandem FP (sctFP) barcodes, involves linking different FPs within a single polypeptide chain (**Fig. 1A**). We demonstrate that sctFP barcodes can be reliably distinguished by analyzing the fluorescence signal intensity at different wavelengths. This method also supports a simple pixelwise image analysis protocol that is robust and does not require prior segmentation of individual cells, thus enabling automated image processing. Unlike previous barcoding strategies limited to certain types of biosensors, the defined emission ratios of sctFP barcodes accommodate a broader range of biosensors.

**Figure 1.**
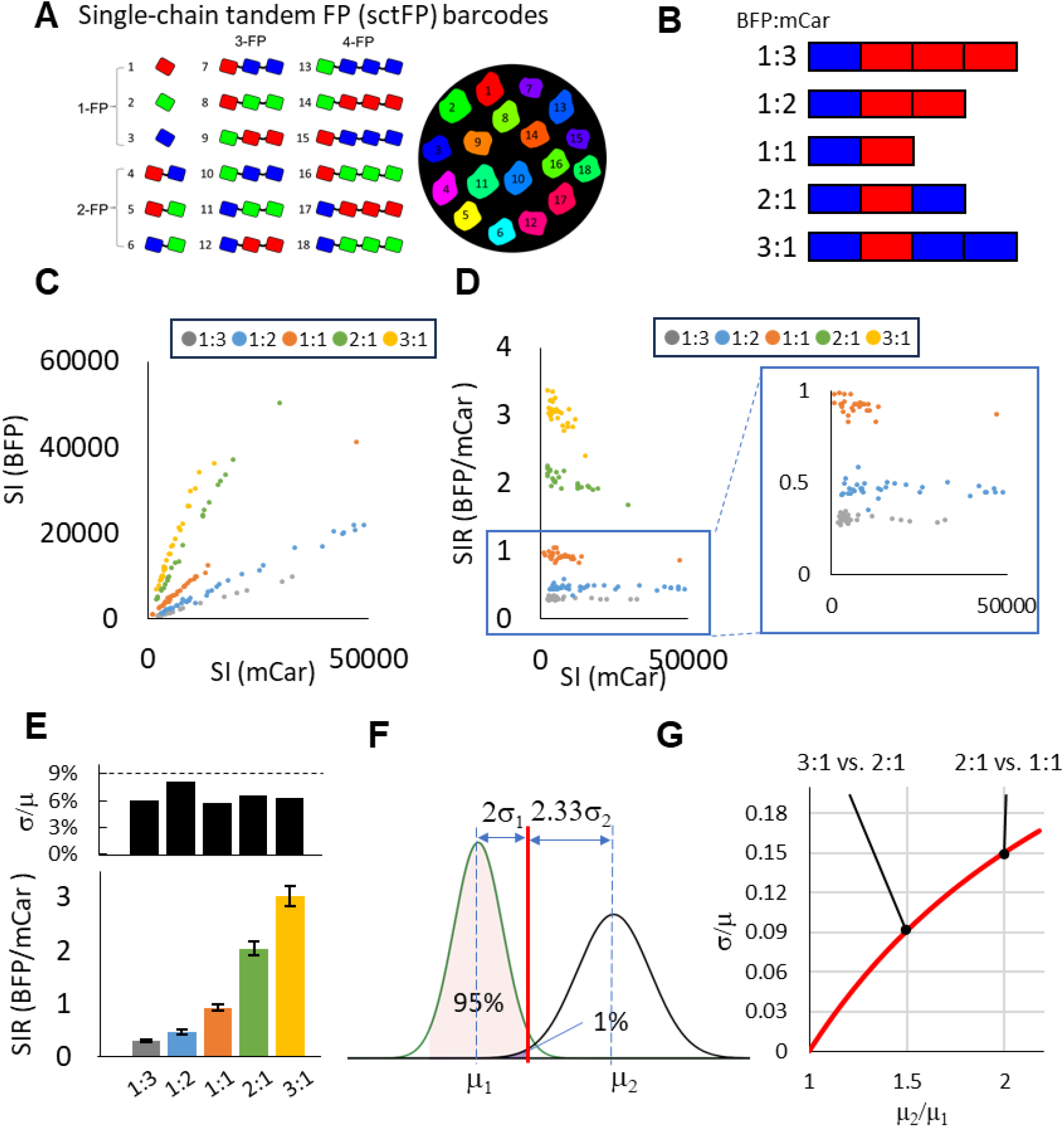
Tandem FP barcodes. (A) sctFP barcodes are generated by fusing FP of different colors at various copy numbers. (B) Barcodes made from five different copy number ratios of BFP and mCardinal. (C) Plot of BFP vs. mCar signal intensity (SI) of individual cells expressing the five sctFP barcodes. (D) Plot of BFP/mCardinal signal intensity ratio (SIR) vs. mCardinal signal intensity of individual cells expressing the five barcodes. (E) The BFP/mCardinal SIR (mean ± SD of n=29, 35, 24, 27, and 28 cells) for the five barcodes. The SD/mean (σ/μ) of BFP/mCar SIR for each barcode is shown on the top. (F) Hypothetical distribution of the SIR in cells expressing two sctFP barcodes, modeled as normal distributions with means ± SD (standard deviation) of μ_1_ ± σ_1_ and μ_2_ ± σ_2_, respectively. The red line indicates a threshold that includes 95% of barcode 1 while excluding all but 1% of barcode 2. (G) The maximum allowable σ/μ that satisfies the condition in (F), plotted as a function of the μ_2_/μ_1_ ratio.

## Results

### A new barcoding platform based on single-chain tandem fluorescent proteins

To test the sctFP barcoding strategy, we fused BFP (EBFP2, [23]) with the red FP mCardinal [24] at 1:3, 1:2, 1:1, and 2:1, and 3:1 ratios (**Fig. 1B**). We imaged HeLa cells transfected with these barcodes and plotted the fluorescence signal intensity of BFP and mCardinal of individual cells (**Fig. 1C**). As expected, cells expressing the same barcodes fell on straight lines with slopes reflecting the copy number ratios of BFP and mCardinal (**Fig. 1C**). Importantly, the BFP/mCardinal signal intensity ratio (SIR) for individual cells were separated into five non-overlapping groups regardless of the expression level (**Fig. 1D**). Between the groups, the relative SIR of the five barcodes closely matched the expected ratios based on the copy number of BFP and mCardinal (**Fig. 1E**). These results establish the proof-of-principle that the SIR between FPs can be reliably used to distinguish different sctFP barcodes.

To quantitatively understand how SIR variations affect the ability to distinguish between sctFP barcodes, we considered a hypothetical scenario involving sctFPs composed of two FPs, FP1 and FP2, at two different copy number ratios. Assuming a normal distribution of the SIR for the two sctFPs with mean ± SD (standard deviation) = μ_1_ ± σ_1_ and μ_2_ ± σ_2_, respectively (*μ*2 > *μ*1), we estimated the SD required to achieve a minimum 95% identification rate of the two sctFPs in a 1:1 mixture with no more than a 1% error rate (**Fig. 1F**). In this scenario, the threshold for distinguishing barcode 1 (**Fig. 1F**, red line) need to be at least 2σ_1_ from μ1 and at least 2.33σ_2_ from μ2, i.e.

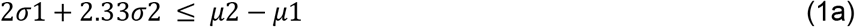

Similarly, the threshold also needs to be at least 2.33σ_1_ from μ1 and at least 2σ_2_ from μ2, i.e.

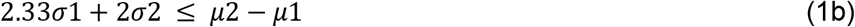

Since the same pair of FPs is used for sctFP construction, it is reasonable to assume that the SD scales with the mean, i.e. σ_1_/μ_1_=σ_2_/μ_2_. This is supported by observing the σ/μ (SD/mean) of the SIR for the five BFP-mCardinal sctFP barcodes (**Fig. 1F**, top). Denoting the σ/μ ratio as *y*, we have σ_1_ = μ_1_ *y* and σ_2_ = μ_2_ *y*. Substituting and rearranging (1) and (2) give

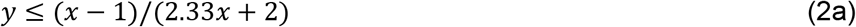

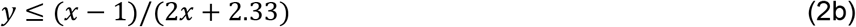

where *x* = *μ*2/*μ*1 is the ratio between the mean SIR of the two sctFP constructs. Note that inequality (2a) is more stringent than (2b) as *x* > 1, and gives the upper limit for σ/μ that enables reliable separation of two sctFP constructs with known FP1/FP2 ratios. For example, distinguishing 2:1 and 1:1 barcodes (corresponding to μ_2_/μ_1_ = 2) requires σ/μ to be smaller than 15%, whereas distinguish 3:1 and 2:1 barcodes (corresponding to μ_2_/μ_1_ = 1.5) requires σ/μ to be smaller than 9% (**Fig. 1G**). Note that the σ/μ for the BFP-mCardinal constructs was below 9% (**Fig. 1E**), thus enabling the separation of all five sctFP constructs in **Fig. 1B**.

### Expanding sctFP barcodes

We next sought to expand the sctFP barcode repertoire by incorporating additional FP combinations. This would enable a broader range of barcodes suited for diverse applications with varying constraints on barcode color selection. Based on the above analysis, we screened FP pairs to identify those with an SIR σ/μ below 15%, which would allow the generation of at least three distinct sctFPs (at 1:1, 1:2, and 2:1 ratios).

Using a selected panel of FPs, we generated sctFPs composed of one copy each of two different FPs (**Fig. 2A, Fig. S1**). We measured their fluorescence signals across multiple wavelengths (**Fig. 2B, Fig. S2A**). The resulting spectral profiles were generally consistent with predictions based on FPbase data [25] (**Fig. 2C,D; Fig. S2**). We calculated SIRs using the channels with peak intensities for the two component FPs (**Fig. 2E**) and determined their corresponding σ/μ values (**Fig. 2F**). Surprisingly, not all sctFPs exhibited consistent SIRs across the cell population, with some FP pairs showing σ/μ values exceeding the 15% threshold (**Fig. 2D,E**; **Fig. S2**).

**Figure 2.**
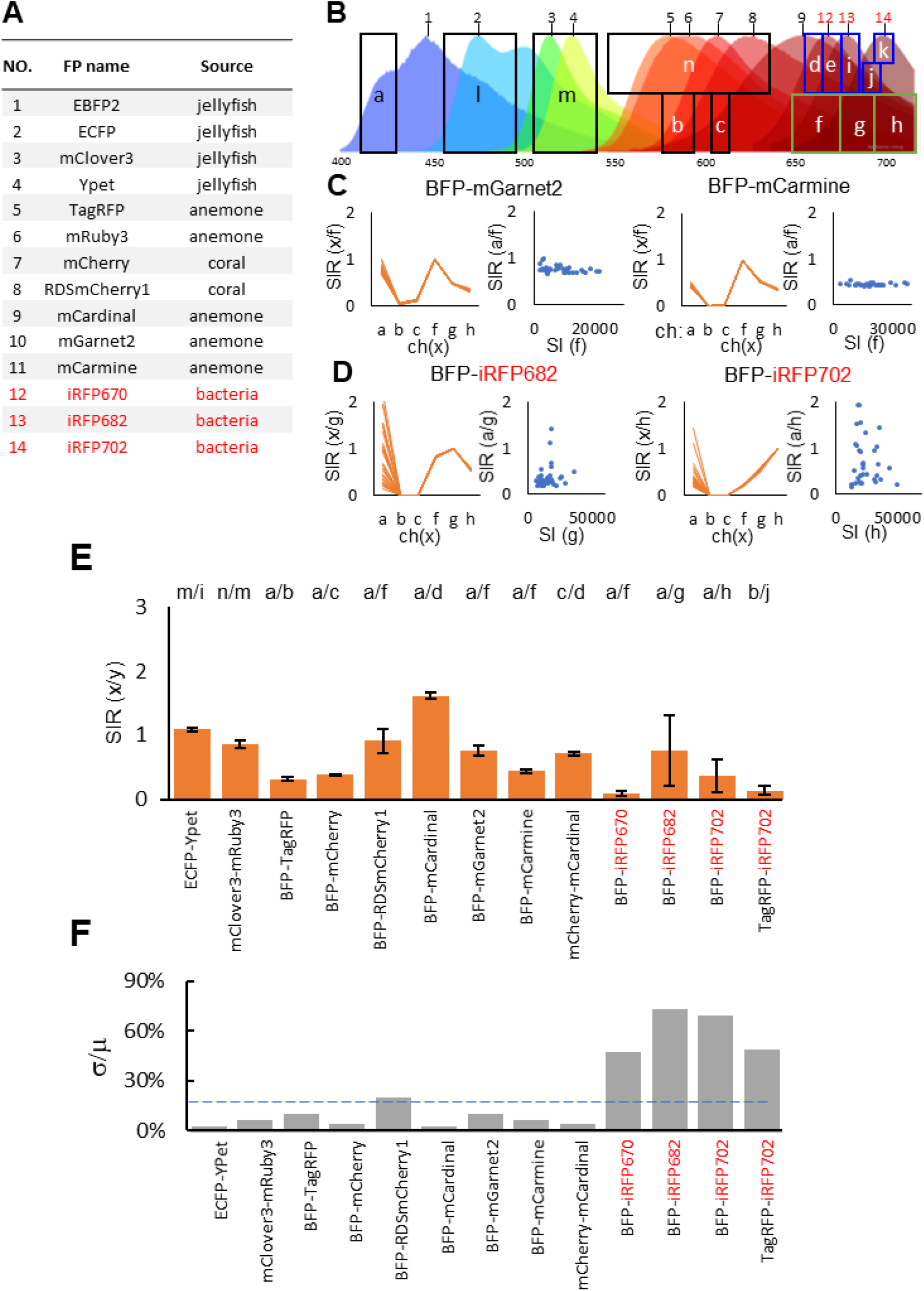
Expanding sctFP barcodes. (A) FPs and their sources. (B) Emission spectra of the FPs shown in (A), along with the acquisition ranges used for channels. The spectra are generated using the spectra viewer from FPbase [25]. Spectra of 10 (mGarnet2) and 11 (mCarmine) are not shown due to overlapping with 12 (iRFP670). (C-D) Emission profiles represented by the SIR of the indicated channels (orange, left) in individual HeLa cells expressing two BFP-RFP sctFP barcodes that showed consistent (C) and variable (D) BFP/RFP SIR (blue dots, right). (E-F) SIR (E) and σ/μ (F) of various sctFP constructs generated from combinations of FPs in (A). Prokaryotic FPs are shown in red.

We selected three FP pairs with σ/μ values below 15%, including BFP-TagRFP, BFP-mCherry, and BFP-mCardinal, for further evaluation of their robustness in mixed populations. These FP pairs were previously used to barcode cells expressing CFP-, GFP-, and YFP-based biosensors due to their spectral separation from the biosensor emission ranges [19] (**Fig. 3A**). Spectral profiles from individual cells expressing each of the three barcodes showed distinct, highly conserved patterns across the population, enabling reliable FP identification (**Fig. 3B**). Importantly, their SIR remained consistent when expressed in different cell lines (**Fig. 3C**).

**Figure 3.**
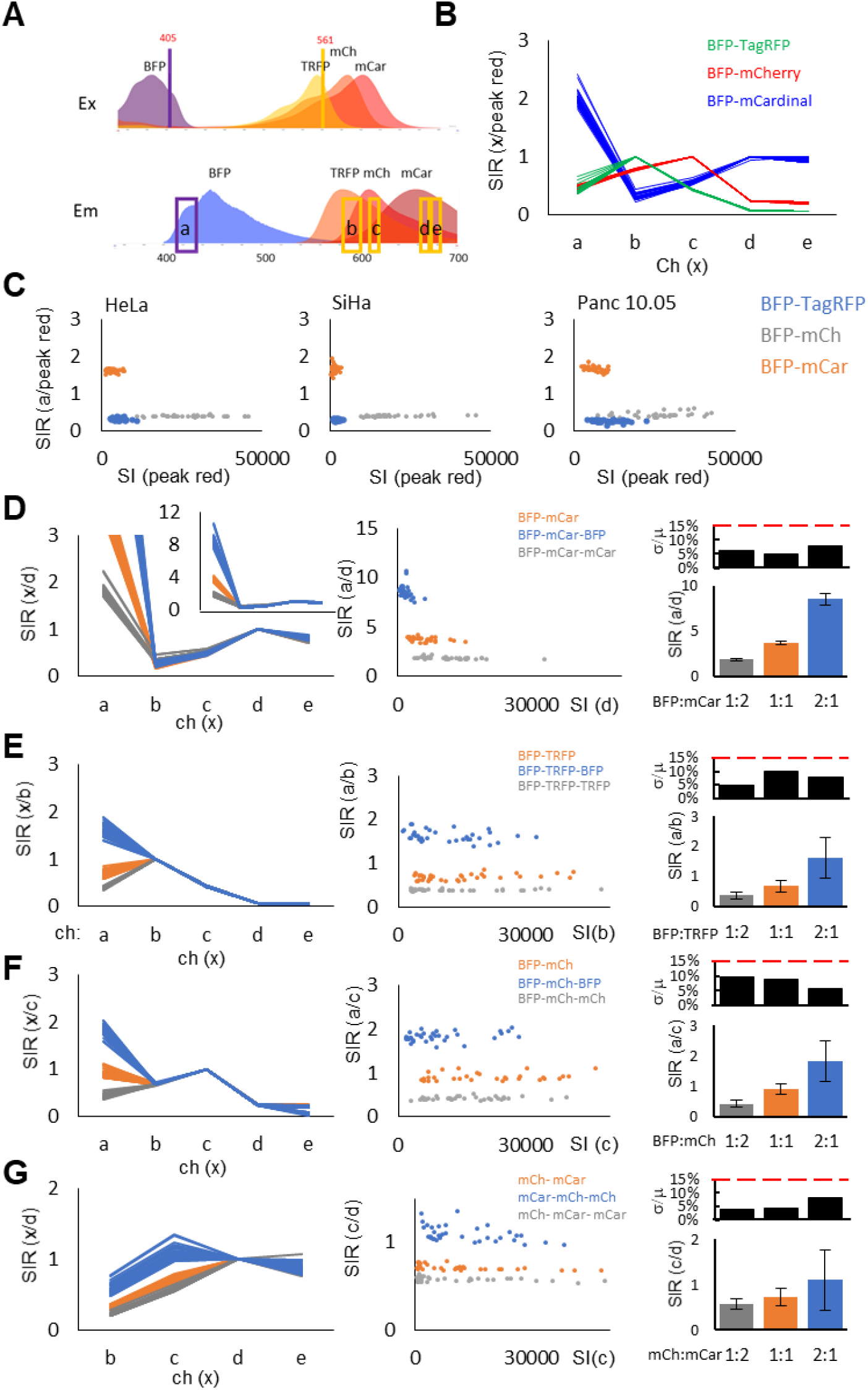
Tandem FP barcodes based on BFP and three red FPs. (A) Excitation and emission spectra of BFP (EBFP2) and three red FPs (RFPs), as well as the excitation wavelengths (vertical lines) and the emission ranges (boxes) for image acquisition. The spectra are generated using the spectra viewer from FPbase [25]. (B) Emission profile of sctFP barcodes made from three BFP-RFP pairs at 1:1 ratio. (C) SIR (BFP/peak RFP) the three sctFP barcodes in (B) expressed in three different cell lines. (D-G) Emission profile and SIR of indicated channels for individual HeLa cells expressing sctFP barcodes made from BFP-mCardinal (D), BFP-TagRFP (E), and BFP-mCherry (F), and mCherry-mCardinal at 1:2, 1:1, and 2:1 ratios. Bar graphs on the right show the mean ± SD as well as SD/mean (σ/μ) of SIR of n=35 cells.

We next imaged cells expressing sctFP barcodes generated by fusing BFP with TagRFP, mCherry, or mCardinal at 1:1, 1:2, and 2:1 ratios. As expected, barcodes incorporating the same red FP exhibited identical spectral profiles within the red emission range (**Fig. 3D-F**, left). Moreover, the BFP/RFP SIR scaled proportionally with the copy number ratios (**Fig. 3D-F**, middle, right). Consistent with their low σ/μ values, the distributions for each barcode were clearly separated with no overlap between groups (**Fig. 3D-F**, middle). Together, these results demonstrate that FP pairs meeting the 15% σ/μ threshold can reliably generate three distinguishable sctFP barcodes based on their spectral profiles and SIRs.

In addition to BFP-red FP combinations, we tested barcodes composed solely of red FPs by fusing mCardinal and mCherry at 1:1, 1:2, and 2:1 ratios. Due to the overlapping spectrum of mCardinal and mCherry, the SIR did not scale proportionally with their copy number ratios. Nevertheless, these barcodes displayed distinct spectral profiles and SIRs, enabling robust classification (**Fig. 3G**).

### FPs that require exogenous cofactors show variable fluorescence intensity ratios

Our analysis revealed high SIR variability when one of the FPs belonged to the iRFP family, which consists of near-infrared (NIR) FPs derived from prokaryotes (**Fig. 2D-F**). To investigate the basis of this variability, we examined images of cells expressing BFP-iRFP constructs. Interestingly, across the population, iRFP signals appeared more uniform than BFP signals (**Fig. 4A**). When we plotted iRFP versus BFP fluorescence, we observed a proportional relationship at low expression levels, but iRFP signal plateaued at higher BFP levels (**Fig. 4B**). For comparison, BFP and mCardinal signals fell on a straight line across the entire range (**Fig. 1C**).

**Figure 4.**
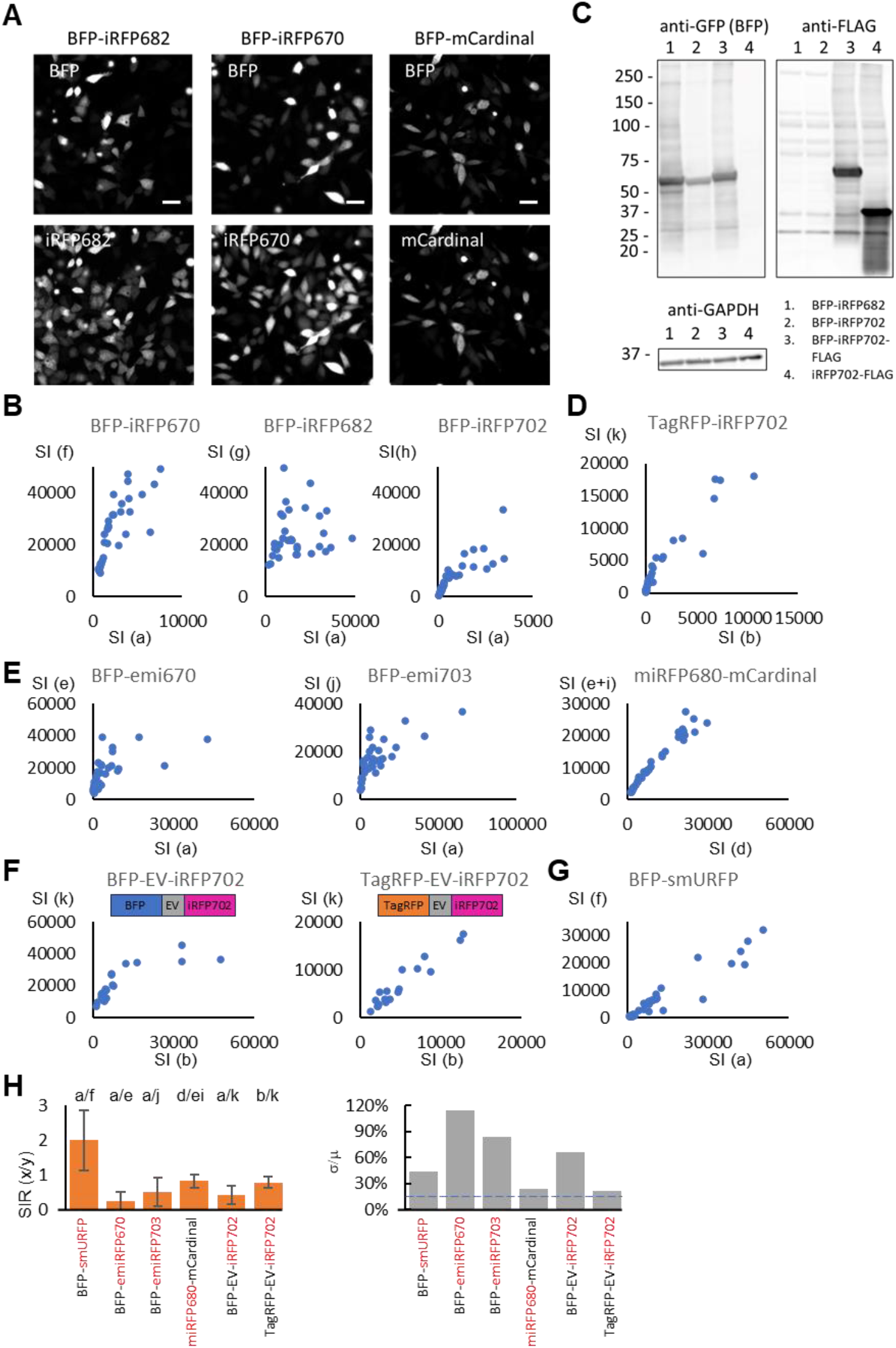
FPs that require exogenous cofactors show variable fluorescence signal intensity. (A) Images of HeLa cells expressing sctFP barcodes made from BFP-iRFP at 1:1 ratio. (B) Plots of FP signal intensity (SI) (see **Fig. 2A** for channel nomenclature) of individual cells expressing the three BFP-iRFP sctFP barcodes. (C) Immunoblots for BFP (anti-GFP), FLAG, and GAPDH of various BFP-iRFP and the iRFP constructs. (D-G) Plots of FP signal intensity (SI) (see **Fig. 2A** for channel nomenclature) of individual cells expressing sctFP barcodes containing TagRFP-iRFP702 (D), monomeric iRFPs (E), the EV-linker (F), and smURFP (G). (H) Bar graphs summarizing the SIR (mean ± SD of n=35 cells) and SD/mean (σ/μ) for the various constructs. The dotted line indicates the 15% σ/μ threshold.

To rule out protein degradation, we performed immunoblotting. Since BFP is derived from GFP, we used an anti-GFP antibody to detect BFP, and added a FLAG tag to iRFP for detection with an anti-FLAG antibody. Immunoblotting analysis confirmed that both FPs were expressed primarily as full-length fusion proteins (**Fig. 4C**). The variability was not specific to the BFP-iRFP combination, as constructs using TagRFP-iRFP702 also showed high SIR variability across the population (**Fig. 4D**).

To rule out the possibility that iRFP dimerization caused spectral interference, we generated sctFPs using monomeric iRFP variants—including emiRFP703, emiRFP670, and miRFP680 [26]—fused to BFP or mCardinal. None of these constructs showed consistent SIRs (**Fig. 4E**). We further tested whether intramolecular FP interactions contributed to the variability by inserting an EV linker, a long, flexible peptide designed to minimize baseline interactions between FPs in biosensor constructs [27]. However, both BFP–EV– iRFP702 and TagRFP–EV–iRFP702 still displayed high SIR variability (**Fig. 4F**).

These findings suggest that iRFP fluorescence intensity does not scale linearly with protein abundance. A likely explanation is that, unlike cnidarian FPs, iRFPs require incorporation of biliverdin as a chromophore [28,29]. Biliverdin incorporation rate may vary between cells, particularly at high expression levels where it becomes limiting. Consistent with this idea, previous studies have shown that exogenously added biliverdin enhances iRFP fluorescence [30].

Together, our results indicate that cnidarian-derived FPs—such as BFP and YFP (jellyfish), mCherry (coral), and TagRFP (anemone)—exhibit robust and reproducible SIRs because they do not rely on exogenous chromophores. In contrast, FPs that require external factors like biliverdin show greater variability. To further test this hypothesis, we evaluated BFP–smURFP constructs, as smURFP is a cyanobacteria-derived FP that also requires biliverdin [31]. As expected, BFP–smURFP barcodes exhibited high SIR variability across the population (**Fig. 4G**). The SIR and σ/μ for various constructs are summarized in **Fig. 4H**. It should be noted that while BFP–iRFP constructs at varying ratios cannot be reliably distinguished based on SIR, they remain distinguishable from BFP-only or iRFP-only barcodes (**Fig. S3**).

### sctFP barcodes enable high-throughput image analysis of biosensor activities

We applied sctFPs to enable multiplexed tracking of signaling activities in live cells using genetically encoded fluorescent biosensors. In this strategy, different populations of cells are separately transfected with a distinct biosensor paired with a unique sctFP barcode. The cells are then pooled and imaged together under the same perturbation conditions. Signals from cells expressing the same biosensor are grouped by barcode to reconstruct the corresponding signaling dynamics over time (**Fig. 5A**).

**Figure 5.**
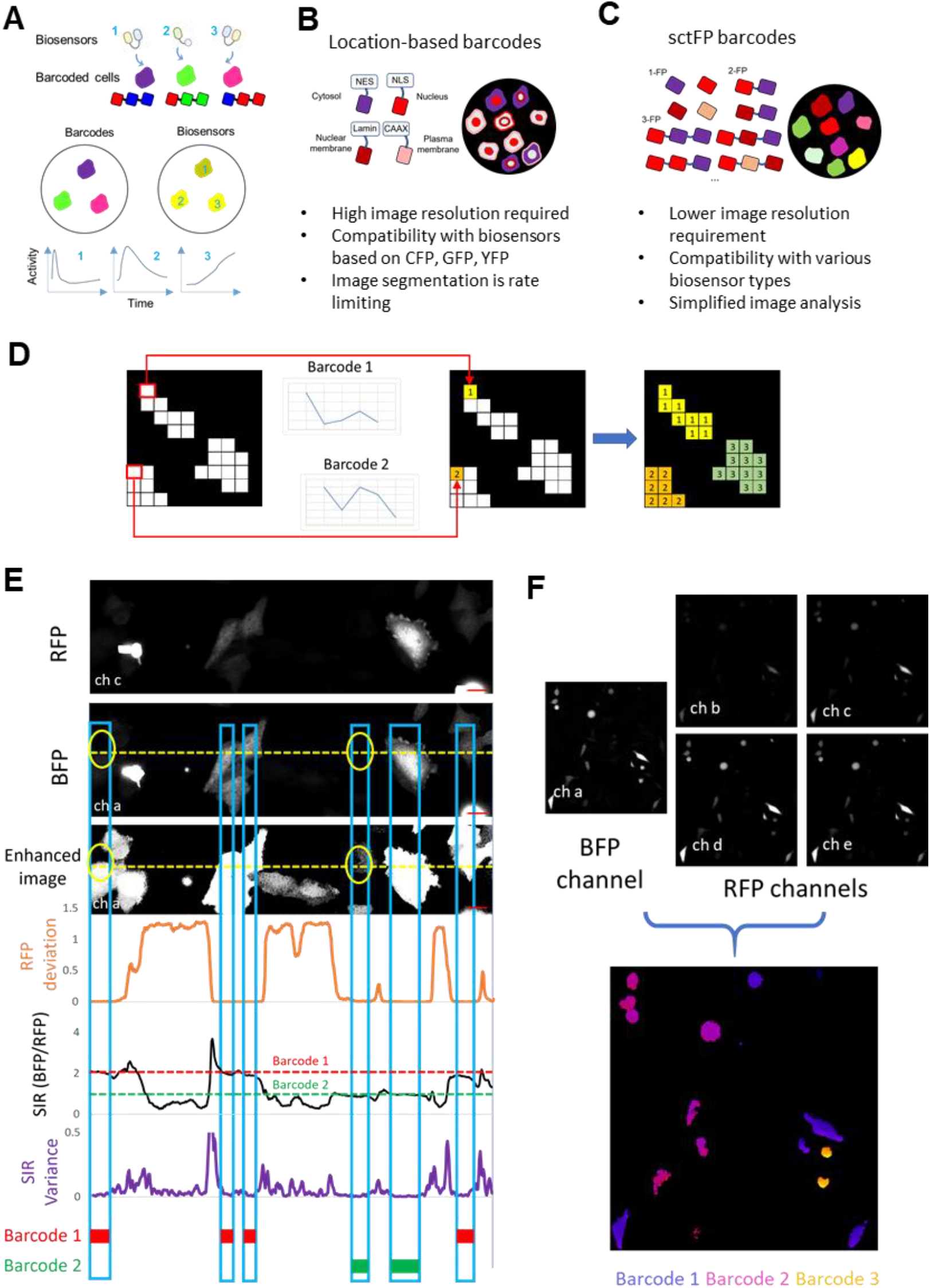
sctFP barcodes facilitate automated image analysis. (A) Schematic of multiplexed biosensor imaging using barcoded cells. Each cell is transfected or transduced with a specific biosensor and a unique barcode. Cells are then mixed for simultaneous imaging, and biosensor activity is extracted from cells based on their corresponding barcodes. (B-C) Comparison of location-based barcodes (B) and sctFP barcodes (C). (D) Schematic of pixelwise analysis enabled by sctFP barcodes, where each pixel is assigned a barcode based on its emission profile. Cell masks are then generated by grouping neighboring pixels sharing the same barcode. (E-F) An example of barcode determination using pixelwise analysis (see text for details).

A previous implementation of this concept used location-based barcodes, in which pairs of fluorescent proteins were targeted to distinct subcellular compartments [19] (**Fig. 5B**). While effective with high-resolution confocal imaging, accurately identifying the subcellular localization of barcode proteins becomes challenging in lower-resolution images typically used for high-throughput screening, particularly in cells with scant cytoplasm. These challenges are often exacerbated by uneven expression of the barcoding proteins. Additionally, directing FPs to different subcellular compartments may differentially affect cell physiology. As discussed below, inconsistent expression levels between the two FPs further limit the applicability of this approach to biosensors outside the CFP, GFP, and YFP emission range. We reasoned that the simpler sctFP design could overcome these limitations by enabling robust barcode identification from lower-resolution images, increasing compatibility with high-throughput imaging, and expanding the usable biosensor spectral range (**Fig. 5C**).

A key advantage of sctFP barcodes is their simplified image analysis. For location-based barcodes, identification of the subcellular localization of FPs starts with segmentation of individual cells [19,20]. Theoretically the process can be accomplished using machine learning models trained on different barcode images. However, due to the heterogeneity of the spatial patterns of barcodes, model training would require a large number of images encompassing the full range of cell density, fluorescence intensity, and spatial patterns, and new models will likely be needed for each cell type due to differences in cell morphologies or imaging parameters. A significant fraction of cells (sometimes over 50%) are also eliminated during analysis due to ambiguous location patterns, as high-magnification imaging is limited by a thin focal plane (< 500 nm) and many cells in the field of view can be out of the focal plane. Therefore, image segmentation has been the rate limiting step in the analysis of biosensor barcoding images [19,20].

sctFP barcodes can circumvent the image segmentation problem: since the barcode can be uniquely determined by the SIR of different FPs, spatial information is not necessary for barcode identification. As a result, sctFP barcodes can be imaged using low magnification objectives, which allow relatively thick focal plane. Then each pixel can be assigned a barcode based on its spectral profile. Masks created for each barcode can then be used to define individual cell areas (**Fig. 5D**). A key benefit of pixelwise analysis is that it eliminates the need to define individual cell boundaries, which are often irregular or poorly defined, particularly for dim cells.

We tested this approach using an image of mixed cells expressing BFP-RFP (mCardinal) sctFP barcodes at 1:1, 1:2, and 2:1 ratios. By analyzing signals across different channels, we computed deviations from a reference RFP intensity profile for pixels along a horizontal line intersecting multiple cells (**Fig. 5E**, orange plot). This deviation plot effectively identified cellular pixels, including those with low signals that only became apparent upon contrast adjustment (**Fig. 5E**, yellow ovals). We also plotted the BFP/RFP intensity ratio along the same line (**Fig. 5E**, black plot). Due to chromatic aberration, caused by wavelength-dependent differences in refraction, the BFP/RFP SIR was skewed near cell edges, increasing pixel-to-pixel variance (**Fig. 5E**, purple plot). By excluding pixels with high SIR variance, we accurately assigned barcodes based on BFP/RFP SIR (**Fig. 5E**, red and green rectangles). Applying this analysis across the full image field yielded regions corresponding to distinct barcodes, which aligned well with cellular boundaries (**Fig. 5F**).

We next used sctFP barcodes to track the responses of RTK network biosensors to epidermal growth factor (EGF) stimulation. We barcoded cells expressing eight FRET-based biosensors and a CFP control using sctFPs composed of BFP paired with TagRFP, mCherry, or mCardinal at 1:1, 1:2, and 2:1 ratios (**Fig. 6A**). The biosensors detect the activity of RTK (EGFR) and its downstream signaling proteins (**Fig. 6B**). Images were acquired across BFP and four red FP channels for barcode identification, and CFP and YFP channels for biosensor readout, following EGF stimulation. Image analysis proceeded as follows: (1) red FP profiles were used to identify TagRFP, mCherry, and mCardinal (**Fig. 6C**, step 1); (2) the BFP/peak red FP ratio was used to assign barcode identities (**Fig. 6C**, step 2); and (3) YFP/CFP ratios, used as a proxy for biosensor FRET level, were calculated pixelwise and averaged across pixels sharing the same barcode (**Fig. 6C**, step 3). We plotted both raw YFP/CFP values (**Fig. 6D**) and values normalized to prestimulus levels (**Fig. 6E**). The biosensor responses were consistent with previous reports of EGF-stimulated signaling [19], while the CFP control showed no response, as expected (**Fig. 6D**,**E**). Importantly, the entire analysis was fully automated. These results demonstrate that sctFP barcoding facilitates high-throughput image analysis by obviating the need for cell segmentation.

**Figure 6.**
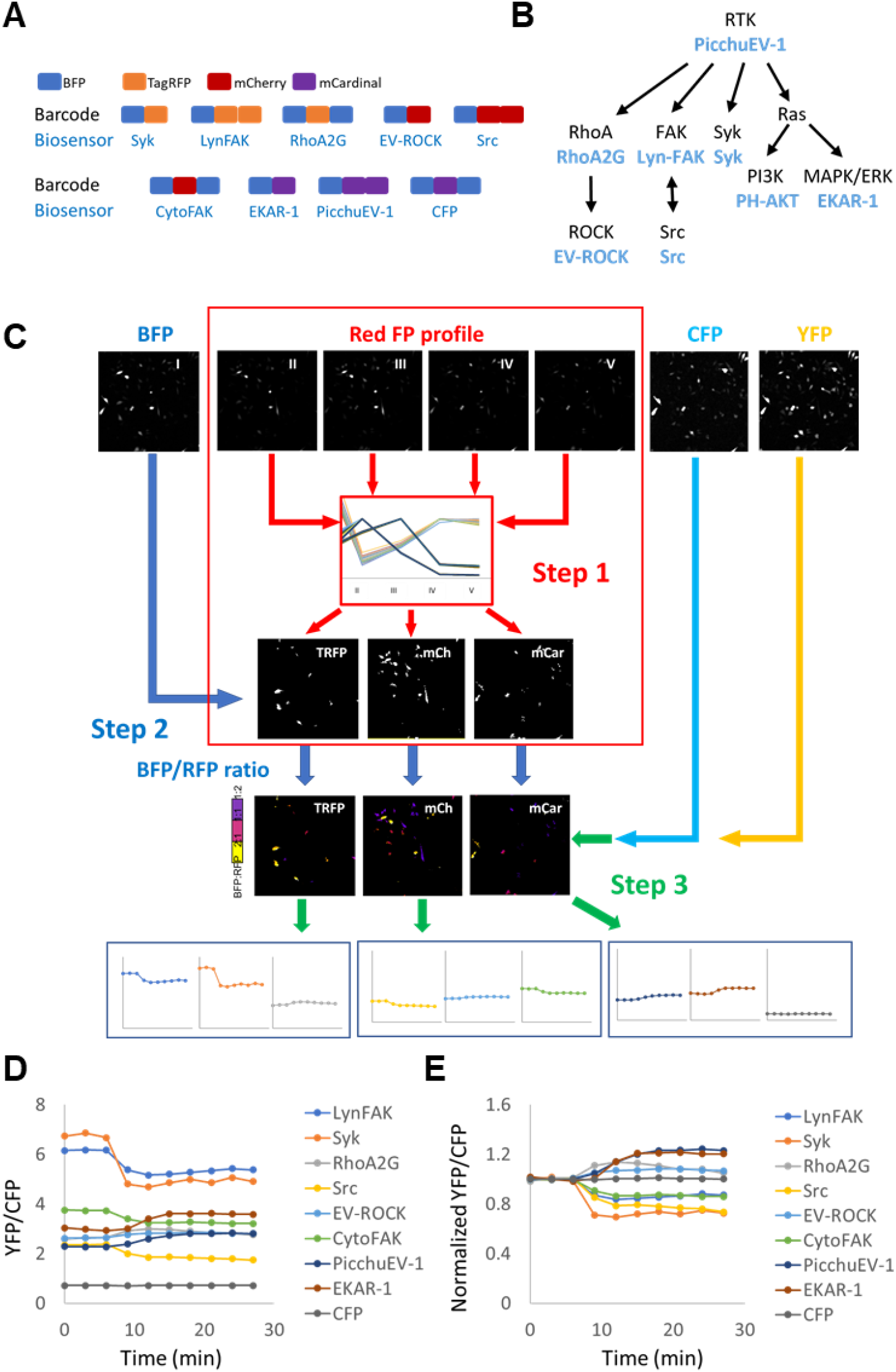
Multiplexed tracking of biosensor activities using sctFP barcodes. (A) sctFP barcodes made from BFP and three RFPs (TagRFP, mCherry, mCardinal) at different ratios were used to label cells expressing 8 CFP-YFP FRET biosensors for the RTK signaling network and CFP (control). (B) Targets of the RTK signaling network reported by the biosensors (shown in blue). (C) Workflow for analyzing multiplexed images. For each pixel, the identity of the RFP is determined from the spectral profile to create cell masks for individual RFPs (Step 1). Next, the BFP/RFP ratio is used to create masks for each barcode (Step 2). Finally, the masks are applied to the CFP and YFP images to calculate the YFP/CFP ratio for each barcode. (D) YFP/CFP ratios for 8 biosensors and CFP (control) in HeLa cells stimulated with 100 ng/ml EGF at 6 min. (E) The YFP/CFP ratios from (C) normalized to prestimulation levels.

### sctFP barcodes enable multiplexed tracking of different types of biosensors

We tested whether sctFP barcodes could be robustly implemented in multiplexed dynamic measurement using biosensors with different spectral properties. Previous barcodes based on BFP and RFPs are spectrally separable from biosensors traditionally made from CFP, GFP, or YFP [19] (**Fig. 5B**). In principle, biosensors and barcodes only need to be distinguishable within the same cells. However, due to the uncoordinated expression level of different FPs in earlier barcode designs, a complete separation between biosensor and barcode spectrums at the population level is essential. This is because accommodating biosensors that spectrally overlap with BFP or RFP may require using single-FP barcodes, which are difficult to distinguish from two-FP barcodes with weak expression of one FP. Although CFP-, GFP-, and YFP-based biosensors remain widely used, alternative fluorescent proteins with enhanced dynamic range and sensitivity are increasingly incorporated into biosensor development. We reasoned that the robust SIR offered by sctFP barcodes could overcome the limitations of previous designs, thereby enabling broader compatibility with a wider range of biosensor types.

We considered the following types of biosensors that involve FPs beyond the CFP-GFP-YFP emission range: 1) GFP-RFP FRET-based biosensors, such as those using the mClover-mRuby3 pair, which offer greater spectral separation, reduced autofluorescence, and lower phototoxicity [32,33]. 2) cpGFP-RFP biosensors, such as the ATP biosensor iATPsnFR [34], which consist of a circularly permuted GFP (cpGFP) whose fluorescence changes upon target binding, fused to a red FP (e.g., mCherry) used for normalization. 3) Excitation-ratiometric (ExRai) biosensors, which typically exhibit high dynamic ranges and use ratiometric measurements based on excitation rather than emission [35–39]. These biosensors are spectrally separable from only a subset of FPs used in the BFP and RFP based barcodes (**Fig. 7A**,**B**). For example, ExRai biosensors require an excitation range that overlaps with BFP and are only compatible with RFP based barcodes. The robust SIR of sctFP barcodes would enable the simultaneous use of BFP-RFP and RFP-only barcodes in the population mix.

**Figure 7.**
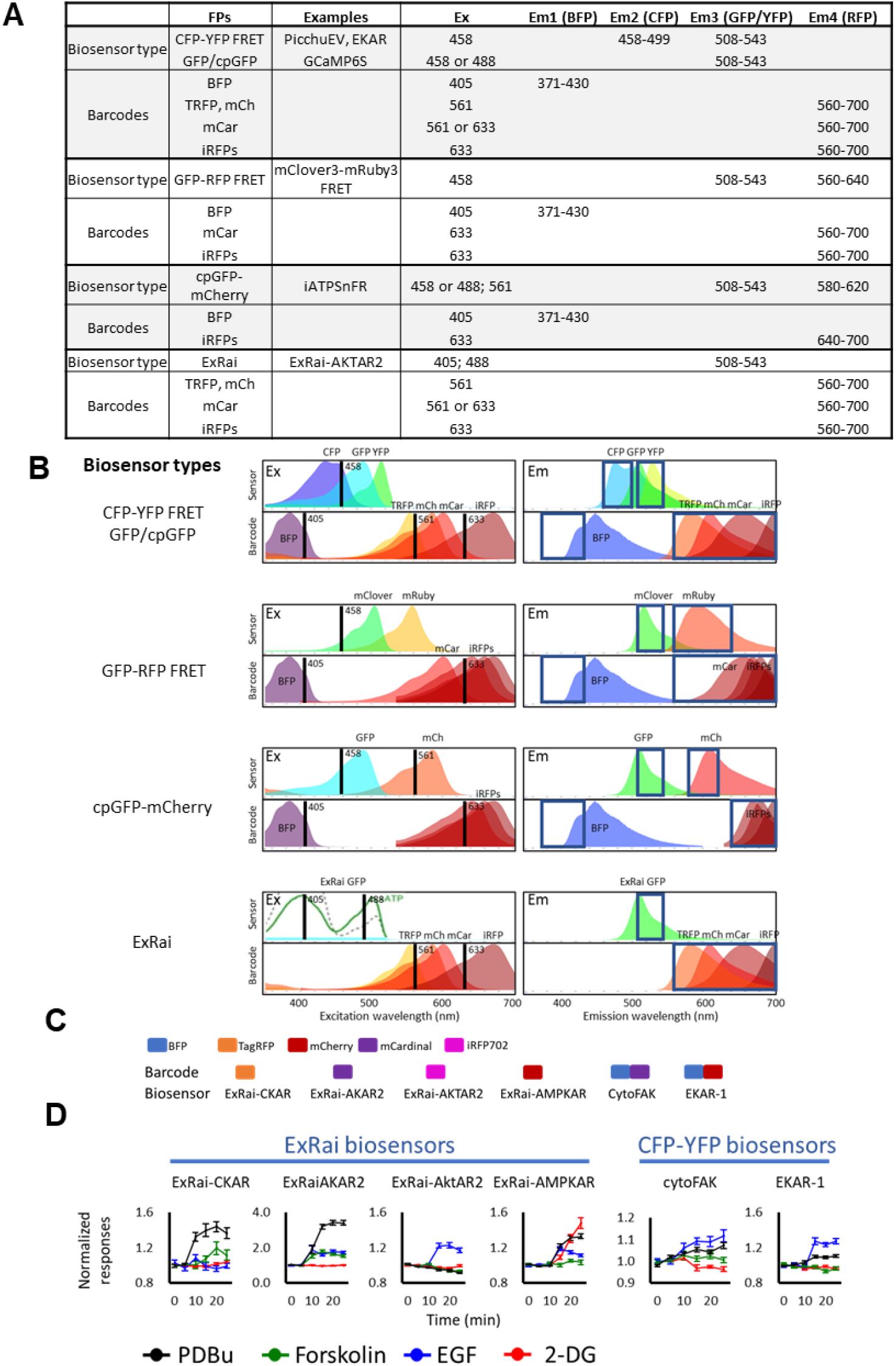
Multiplexed tracking of various types of biosensors. (A) Summary of different types of biosensors and the barcode FPs that are spectrally compatible with each. Shown are the excitation wavelengths and the emission wavelength ranges used for acquiring signals from both biosensors and barcodes. (B) Excitation (Ex) and emission (Em) spectra of FPs used in the biosensors and their compatible barcodes. Vertical black lines indicate the excitation wavelengths, and blue boxes denote the emission ranges used for signal acquisition. (C) sctFP barcodes for ExRai and CFP-YFP FRET biosensors. (D) Simultaneous tracking of ExRai and CFP-YFP FRET biosensors in mixed barcoded cells treated with known activators. The responses (mean ± SEM of n=20 cells) are normalized to prestimulus levels.

As a proof of principle, we tested simultaneous tracking of ExRai and FRET biosensors. We used RFP-only barcodes for cells expressing four ExRai biosensors that detect the activity of PKC (CKAR), cAMP (AKAR), AKT (AKTAR2), and AMPK (AMPKAR) [36,37,40], and BFP-RFP barcodes for cells expressing two CFP-YFP FRET biosensors that detect the activity of FAK (CytoFAK)[41] and ERK (EKAR-1, derived from EKAR [42]) (**Fig. 7C**). Cells were mixed and simultaneously imaged during stimulation with four known activators: 2-deoxy-D-glucose (2-DG) for AMPK, EGF for AKT and FAK, forskolin for cAMP, and phorbol 12,13-dibutyrate (PDBu) for PKC. We were able to reliably distinguish the barcodes and quantify the responses of the corresponding biosensors (**Fig. 7D**). All biosensors responded to their expected stimuli, with magnitudes and kinetics comparable to those observed in pure populations of cells expressing individual biosensors (**Fig. S4**). In addition to the expected responses, we also observed unexpected responses, such as activation of ExRai-AKAR2 by PDBu. The significance of these responses is currently under investigation. Together, these results demonstrate that sctFP barcodes enable simultaneous tracking of biosensors with distinct spectral properties.

## Discussion

In this study, we developed a new method for barcoding cells by combining FPs with distinct spectral properties in varying ratios into a single polypeptide chain. Our findings demonstrate that these sctFP barcodes can be reliably differentiated through the relative intensity of the component FPs. Interestingly, this strategy proved effective for cnidarian FPs, but not for prokaryotic FPs, which exhibited variability in fluorescence signals likely due to their reliance on exogenous cofactors. We also showed that the analysis of sctFP barcodes can be streamlined through a pixelwise analysis approach. Additionally, we successfully applied this method in multiplexed imaging of various types of biosensors designed to detect different signaling activities.

Genetically encoded fluorescent biosensors serve as powerful tools for monitoring signaling dynamics in live cells [43–45]. While these biosensors have primarily been employed to investigate individual molecular activities, there is a growing interest in multiplexing biosensors to gain insights into the interactions among various signaling pathways. Most existing strategies focus on distinguishing biosensors expressed in the same cells based on differences in their spectral, spatial, or dynamic properties [46–48]. More recently, an alternative approach utilizing FP-based barcodes has emerged to label cells expressing different biosensors for simultaneous imaging [16,19–22,49]. This barcoding strategy eliminates interference between different biosensors within the same cells, with any potential adverse effects confined to the cells expressing a specific biosensor. Additionally, the approach is easily scalable to accommodate high multiplexity and can readily integrate existing biosensors. In prior implementations of biosensor barcodes, various FPs targeted to distinct subcellular locations were combined to enhance multiplexity. While this method has proven effective, the use of FPs localized to different regions within the cell can produce variable effects and necessitates high-resolution imaging to accurately identify their subcellular locations. Furthermore, imbalances in expression levels of barcoding FPs may hamper barcode identification in subsets of cells.

Our sctFP barcodes effectively address these challenges. They provide defined and reproducible fluorescence signatures, allowing for the use of relative intensity between FPs as the basis for barcoding. Because the intensity of an FP can be measured more accurately than its subcellular localization, sctFP barcodes offer significantly enhanced robustness compared to location-based sensors. Furthermore, these barcodes can be decoded directly from pixel-level spectral data, eliminating the need for time-consuming cell segmentation and enabling fast, fully automated image analysis. This greater tolerance for variations in image resolution and the streamlined image analysis process facilitate adaptation to high-throughput formats, where lower magnification images are often employed. Additionally, the simplified design of sctFP barcodes enhances compatibility with a wider range of biosensors and may synergize effectively with synthetic biology tools. Although not demonstrated in the current study, sctFP barcodes could also be applicable in bacterial cells, which do not possess the subcellular compartments required for location-based barcoding.

While the primary motivation for this work is the multiplexing of genetically encoded fluorescent biosensors, fluorescent cell barcodes have also been developed for various other applications. For instance, Brainbow employs the Cre-Lox recombination system to randomly express different ratios of FPs in individual neurons, facilitating the tracing of neural connections and the study of brain architecture [16]. However, the random nature of Brainbow limits their utility in labeling defined cell populations, which is essential for biosensor barcoding or genetic screening. Pairing a small set of FPs can generate a large number of barcodes that is compatible with genetic screening and lineage tracing [50], and fluorescent tagging of endogenous proteins has been employed to visualize their spatiotemporal dynamics on a large scale under various perturbation conditions [14,51], but these strategies present challenges when integrating additional fluorescent readouts, such as biosensors. Barcodes based on sequential multicolor immunofluorescence of epitope combinations have been utilized for protein engineering [15], but they require additional steps involving cell fixation for immunofluorescence following live-cell imaging. As a result, this method cannot sustain longitudinal monitoring. Our approach provides a simple and robust alternative to existing barcoding strategies, particularly for applications that demand precise detection of fluorescence readouts, such as biosensor use.

In summary, the sctFP barcodes represent a simple and versatile tool for studying signaling dynamics and beyond.

## Supporting information

Wu et al SOM

## Acknowledgments

This work was supported by funding from R01GM136711 (to C.H.H.), Cervical Cancer SPORE P50CA098252 Career Development Award (to J.M.Y.) and Pilot Project Award (to C.H.H.), the Sol Goldman Pancreatic Cancer Research Center (to C.H.H.), and the Johns Hopkins Discovery Award (to C.H.H.). Additionally, the purchase of the Zeiss LSM 780 confocal microscope by Johns Hopkins University School of Medicine Microscope Facility was made possible through NIH grant S10OD016374.

## Methods

### Cells

HeLa, HEK293T, SiHa and Panc 10.05 cells, purchased from ATCC, were grown at 37°C, 5% CO_2_ in DMEM high glucose medium (Gibco, #11965092) supplemented with 10% FBS (Corning Cellgro, 35-010-CV), 1 mM sodium pyruvate (Gibco, #11360070), and 1X nonessential amino acids (Gibco, #11140076).

### Chemical reagents

Stocks of 200 μM phorbol-12,13-dibutyrate (PDBu, EMD Millipore, #524390),10mM Forskolin (Torcis Bioscience, #1099) were prepared by dissolving the chemicals in DMSO. Stocks were diluted to the indicated final concentrations in the culture medium. The EGF stock solution was prepared by dissolving EGF (Sigma-Aldrich, E9644) in 10 mM acetic acid to a final concentration of 1 mg/ml. All drug stocks were stored at -20°C. 2-Deoxyglucose (2-DG, MilliporeSigma, #D8375) was dissolved in culture medium to 100 mM and used immediately.

### Plasmids

#### sctFP barcodes

To construct the entry vector backbone containing a multiple cloning site (MCS) for tandem barcode gene cloning (**Figure S4A**), a gBlocks gene fragment (5’-ggatccGGCTCTTCCactagtGGAAGCggtaccTGATCTTCTGgagctcTGGGAGGaagcttGTGAGGCAGCAGCGgaattcTGGGGTCCTCCcatatgTGAGGGTCTTCCgcggccgcAAC-3’) was synthesized by Integrated DNA Technologies and first amplified by PCR. The purified product was then re-amplified using primers containing attB sequences to add attB1 site to 5’ and attB2 site to 3’ ends. This second PCR product was purified and recombined with pDONR221 (Invitrogen, #12536017) in a Gateway BP reaction using BP Clonase II enzyme mix (Invitrogen, #11789020), resulting in the entry vector pENTR221-sctFP-MCS.

pENTR221-sctFP-MCS serves as a versatile cloning vector for inserting up to four fluorescent proteins (FPs) in tandem. The designated restriction sites for FP insertion are: FP1 – BamHI and SpeI; FP2 – KpnI and SacI; FP3 – HindIII and EcoRI; FP4 – NdeI and NotI. A Kozak sequence and an ATG start codon must be included at the 5’ end of FP1 after the BamHI restriction site. To maintain the correct reading frame, a G nucleotide should be added to the 3’ end of FP2 and to both the 5’ and 3’ ends of FP3 if it is included. Internal stop codons are present for FP1, FP2, and FP3 when in-frame, while FP4 must include a stop codon at its 3’ end.

Single FP constructs were obtained from Addgene (**Figure S4B**), amplified to add appropriate restriction sites, and then cloned into pENTR221-sctFP-MCS. These various entry vectors were then recombined with the lentiviral destination vector pLex307 (Addgene, #41392) using Gateway LR Clonase II (Invitrogen, #11791020). The resulting expression plasmids were used for both transient transfection and lentivirus production.

To construct the pcDNA3.1-BFP–EV–iRFP702 and pcDNA3.1-TagRFP–EV–iRFP702 plasmids, we first generated the pcDNA3.1-PicchuEV-1 construct by inserting the PicchuEV sequence [52] into pcDNA3.1(+) using the HindIII and BamHI restriction sites. BFP or TagRFP was then inserted by replacing the smaller fragment between HindIII and KpnI, while iRFP702 was introduced by replacing the smaller fragment between BspEI and BamHI.

#### Biosensors

Plasmids for the following biosensors were purchased from Addgene: CytoFAK (Addgene #78300), LynFAK (Addgene #78299), [41], RhoA2G (Addgene #40176) [53], Src (Addgene #78302) [54], Syk (Addgene #125729) [55], EKAR (Addgene #18679) [42], ExRai-CKAR (Addgene #118409), ExRai-AKAR2 (Addgene #161753), ExRai-AKTAR2 (Addgene #184047), and ExRai-AMPKAR (Addgene #192446). PicchuEV and EV-ROCK were generously provided by Dr. Michiyuki Matsuda’s laboratory.

### Cell transfection and lentiviral transduction

Cells were transiently transfected using the GenJet™ In Vitro DNA Transfection Reagent (Ver. II) following the manufacturer’s protocol (SignaGen Laboratories, #SL100489). Briefly, for one 3.5 cm culture dish, 4×10^5^ cells were seeded the day before transfection. On the day of transfection, the culture medium was replaced with 1 mL fresh complete medium without antibiotics. The GenJet-DNA complex was prepared by diluting (1) 1 μg of DNA in 50 μL serum-free DMEM, and (2) 3 μL GenJet reagent in 50 μL serum-free DMEM in two separate tubes, and then mix the two together by adding GenJet into DNA. The complex was allowed to form at room temperature for 15 min, and then added dropwise onto the cells. Cell expression was visible 24 to 48 h post transfection.

For lentiviral transduction, 4×10^5^ cells were seeded in one 3.5 cm culture dish the day before transduction. The culture medium was then replaced with a complete medium containing lentiviruses and polybrene (final concentration 8 μg/mL). Cells were incubated with the viruses at 37°C, 5% CO_2_ overnight and medium replaced the next day. Cell expression was visible 48 h post transduction.

### Lentivirus production

Lentiviruses were collected from HEK293T cells transfected with a transfer plasmid, a packaging plasmid psPAX2 (Addgene #12260), and an envelope plasmid pMD2.G (Addgene #12259) at the mass ratio of 4:3:1. The culture medium was replaced 5 h after transfection and the conditioned medium was collected 24 h later. More fresh medium was added to the transfected cells and the conditioned medium was collected again after another 24 h. The two batches of conditioned medium containing lentiviruses were pooled together and concentrated 10 times with the Lenti-X Concentrator following the manufacturer’s protocol (Takara Bio, #631231). The concentrated lentiviruses were aliquoted, flash frozen in liquid nitrogen, and stored at -80°C.

### Microscopy

Cells were transfected with plasmids encoding barcode constructs and different biosensor constructs in separate wells. Prior to imaging, cells expressing distinct barcode and biosensor combinations were harvested, pooled, and re-plated onto imaging dishes. Confocal microscopy was performed using a Zeiss LSM 780 or 880 equipped with a spectral detector (for barcode emission profiling) and a motorized stage (for automated acquisition of multiple fields of view).

For the sctFP barcodes, spectral imaging was performed on a Zeiss LSM 780 or 880 confocal microscope equipped with a spectral detector. Each fluorescent protein was excited using its optimal laser line (405, 488, 561, or 633 nm), based on its excitation spectrum. For each sctFP, emission signals were collected across 4–5 detection channels, with wavelength ranges selected according to the emission profile of the corresponding FP (see **Fig. 2B** and **Fig. S2A**).

For CFP–YFP FRET biosensor imaging, cells were excited with a 458 nm laser. Emission signals were collected using two detection windows: 457–500 nm for CFP and 507–544 nm for YFP. FRET activity was quantified as the YFP/CFP emission ratio.

For ExRai biosensor imaging, cells were sequentially excited with 488 nm and 405 nm lasers, and emission signals from both excitations were collected in the 507–544 nm range. ExRai activity was represented by the ratio of emission intensities upon 488 nm versus 405 nm excitation.

Biosensor responses were quantified by calculating pixel-by-pixel intensity differences between time-lapse frames separated by defined intervals. To reduce noise, the 2–3 frames immediately prior to stimulation were averaged to establish a baseline. The percentage change in intensity was then calculated by normalizing each difference to the baseline average.

### Image analysis

Images were processed and analyzed with NIH ImageJ and Fiji [56,57]. The signal intensity ratio (SIR) of individual cells was calculated in Microsoft Excel from the signal intensity (SI) of different channels. Microsoft Excel was also used for statistical analysis and to generate graphs.

Pixelwise analysis of sctFP barcodes comprises the following steps: 1) The SI of all channels (typically a BFP channel plus 4 RFP channels) was measured within a 10×10 pixel area and normalized to the SI of the peak RFP channel. 2) The normalized SI values of the RFP channels were compared to reference RFP spectra to calculate the deviation. 3) The variance of the SIR of BFP/RFP across 5 adjacent pixels was calculated. 4) A pixels was assigned a barcode when it met the following conditions: A) low deviation from the RFP reference spectrum; B) low variance in BFP/RFP across neighboring pixels; C) a BFP/RFP within 10% of the reference ratio, based on the empirical distribution of BFP/RFP values. Thresholds for acceptable RFP deviation and BFP/RFP variance were empirically determined.

### Immunoblotting

The following primary antibodies were used: anti-GFP (Cell Signaling Technology, #2956, 1:1000), anti-GAPDH (Cell Signaling Technology, #2118, 1:2000), and anti-FLAG (Millipore, F7425) for detecting FLAG-tagged sctFPs. The secondary antibody used was Donkey anti-Rabbit Alexa Fluor™ 647 (Thermo Fisher Scientific, A-31573, 1:5000).

Immunoblotting was done as described in our previous publication [58]. Briefly, cells were harvested and lysed in 1X RIPA buffer (Cell Signaling, #9806) containing 1X protease inhibitor cocktail (Roche, #11873580001) and 1X phosphatase inhibitor (Sigma, #P5726). Cell lysates were collected after centrifugation. After mixing with sample buffer and boiling at 95°C for 5 min, proteins were separated by SDS-PAGE using 4–20% Criterion™ TGX™ Precast Midi Protein Gels (Bio-Rad, #5671094), and then transferred to the low fluorescence background PVDF membrane (Millipore, #IPFL00005) in an ice bath at 80V for 1h. Membranes were incubated with primary antibodies overnight followed by incubation with secondary antibodies in 5% BSA in TBST at room temperature for 1h. Images were taken by a Pharos Molecular Imager (BioRad).

