## Supplementary material for "Cell barcoding with tandem fluorescent proteins enables high-throughput signaling dynamics analysis": Wu et al SOM

### Figure S1

A

| FP name | Species | Peak emission (nm) | Addgene | Ref |
|---|---|---|---|---|
| EBFP2 | Aequorea victoria | 448 | 55243 | Michael Davidson |
| ECFP | Aequorea victoria | 477 | 78300 | 41 |
| mClover3 | Aequorea victoria | 518 | 74252 | 33 |
| Ypet | Aequorea victoria | 530 | 78300 | 41 |
| TagRFP | Entacmaea quadricolor | 584 | 99271 | Philipp Keller |
| mRuby3 | Entacmaea quadricolor | 592 | 74252 | 33 |
| mCherry | Discosoma sp. | 587 | 39319 | 59 |
| RDSmCherry1 | Discosoma sp. | 630 | 89987 | 60 |
| mCardinal | Entacmaea quadricolor | 659 | 54590 | 24 |
| mGarnet2 | Entacmaea quadricolor | 671 | 104308 | 61 |
| mCarmine | Entacmaea quadricolor | 675 | 109486 | 62 |
| smURFP | Trichodesmium erythraeum | 670 | 80349 | 31 |
| iRFP670 | Rhodopseudomonas palustris | 670 | 45466 | 29 |
| emiRFP670 | Rhodopseudomonas palustris | 670 | 136556 | 26 |
| miRFP680 | Rhodopseudomonas palustris | 680 | 136557 | 26 |
| iRFP682 | Rhodopseudomonas palustris | 682 | 45459 | 29 |
| iRFP702 | Rhodopseudomonas palustris | 702 | 45456 | 29 |
| emiRFP703 | Rhodopseudomonas palustris | 703 | 136558 | 26 |

B

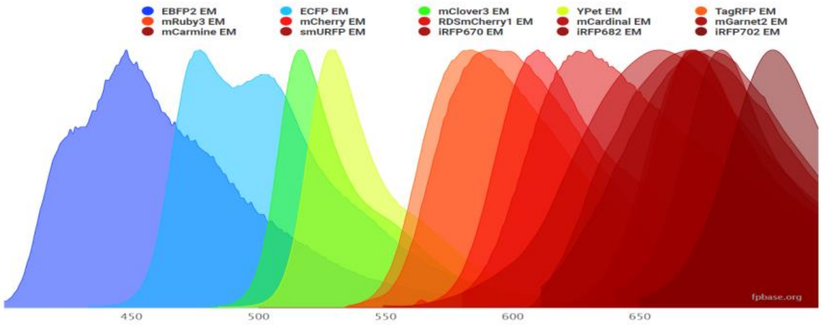


C

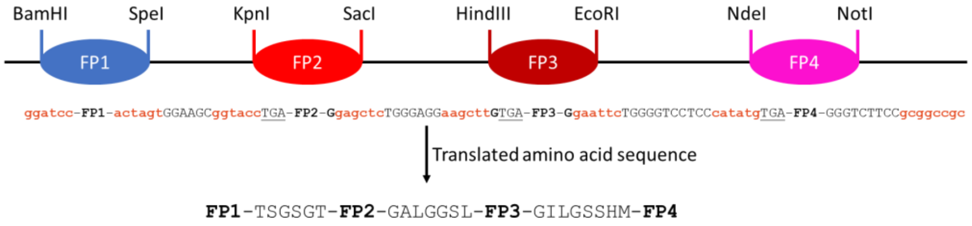


**Cloning notes:**
FP1: Include Kozak sequence and ATG at 5’ end
FP2: add one G at 3’ end
FP3: add one G at 5’ end and another G at 3’ end
FP4: Add STOP codon at 3’ end

**Figure S1. Generation of sctFP barcodes.** (A) FPs used in this study. (B) Emission spectra of the FPs generated using the spectra viewer from FPbase [25]. (C) Schematic of the multiple cloning sites..

### Figure S2

**A**

| Ch. | a | b | c | d | e | f | g |
|---|---|---|---|---|---|---|---|
| Wavelength | 411-431 | 579-597 | 606-615 | 659-668 | 668-677 | 650-677 | 677-695 |
| Ch. | h | i | j | k | l | m | n |
| Wavelength | 697-720 | 677-686 | 691-700 | 697-708 | 458-499 | 508-543 | 550-640 |

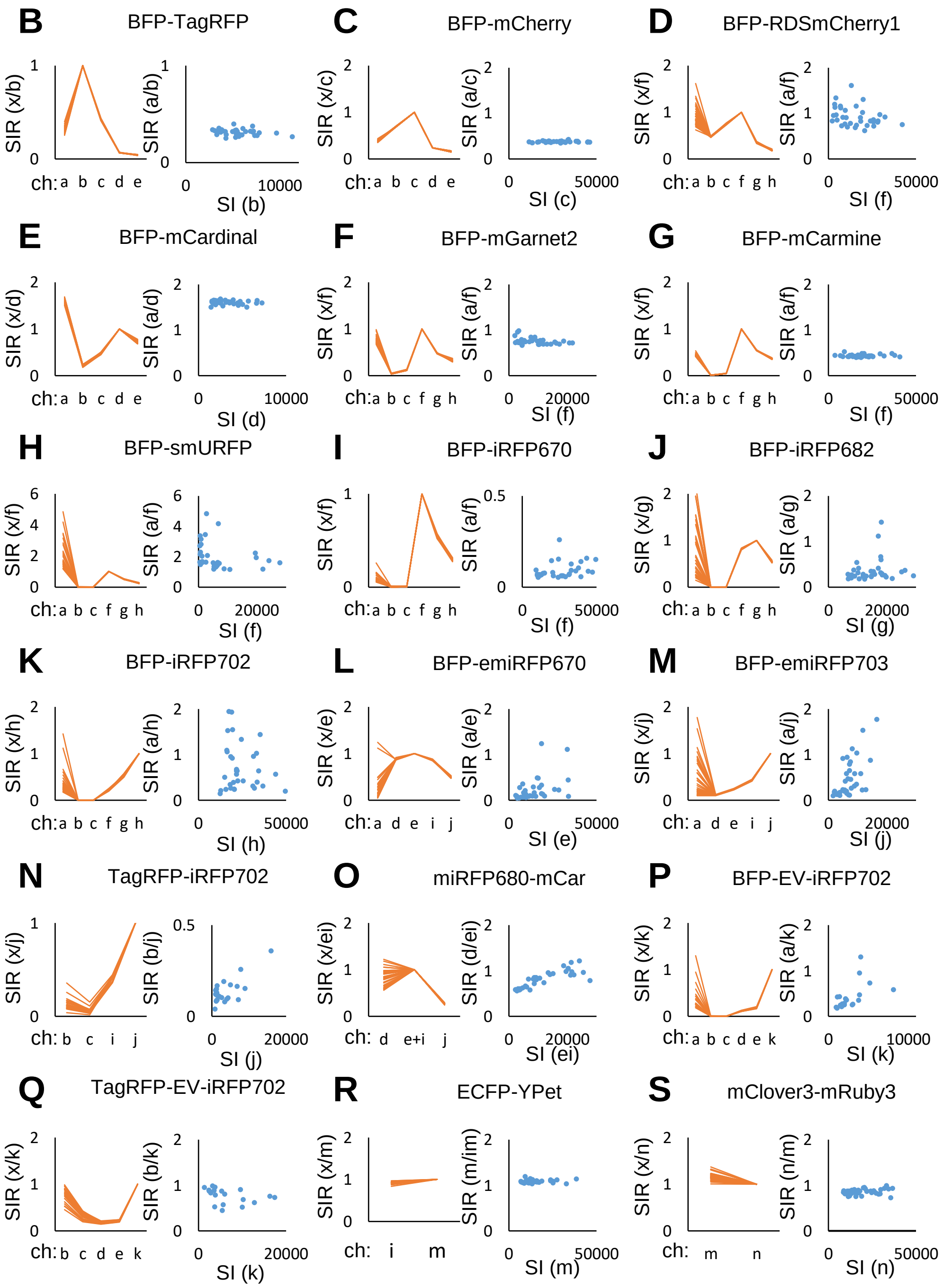


**Figure S2. Characterization of sctFP barcodes composed of single copies of two FPs.** (A) Acquisition ranges of different channels. (B-S) Emission profiles represented by the SIR of the indicated channels (orange, left) and plots of SIR vs. SI of indicated channels (blue dots, right) of individual cells expressing sctFP barcodes.

**Figure S3**

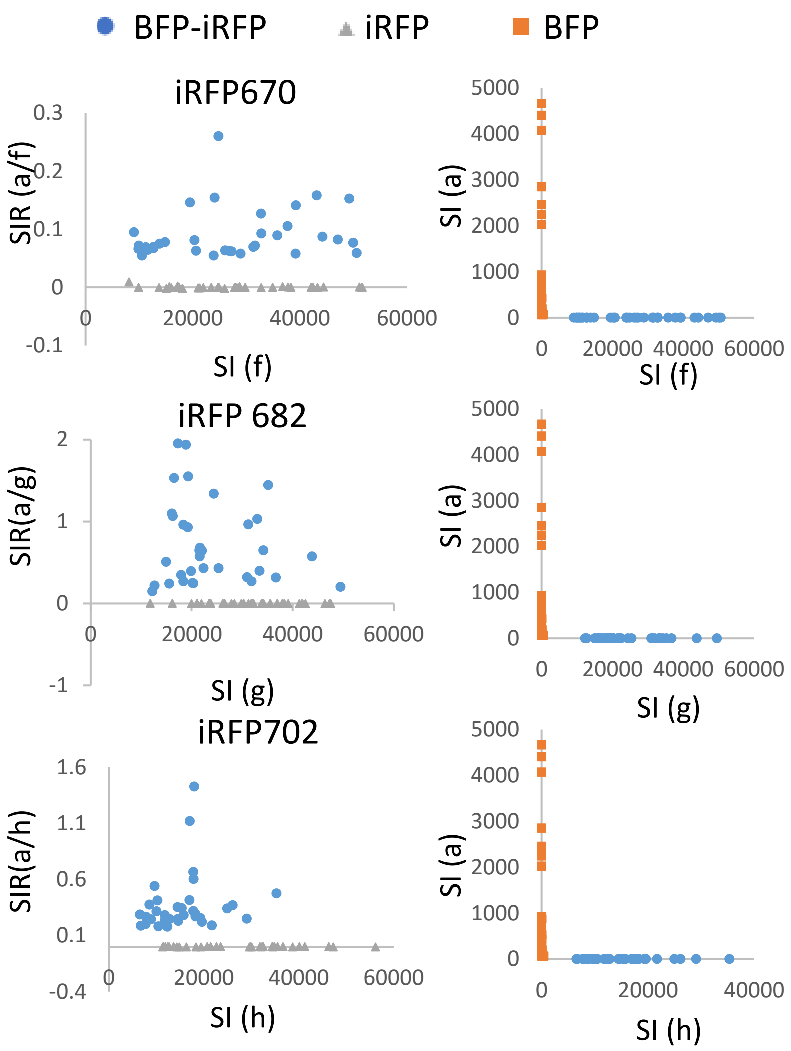


**Figure S3. Comparison of BFP-iRFP sctFP barcodes and individual FPs.** Plots of SIR vs. SI of indicated channels of individual cells expressing sctFP barcodes (blue dots), individual iRFPs (gray triangles), and BFP (orange squares).

### Figure S4

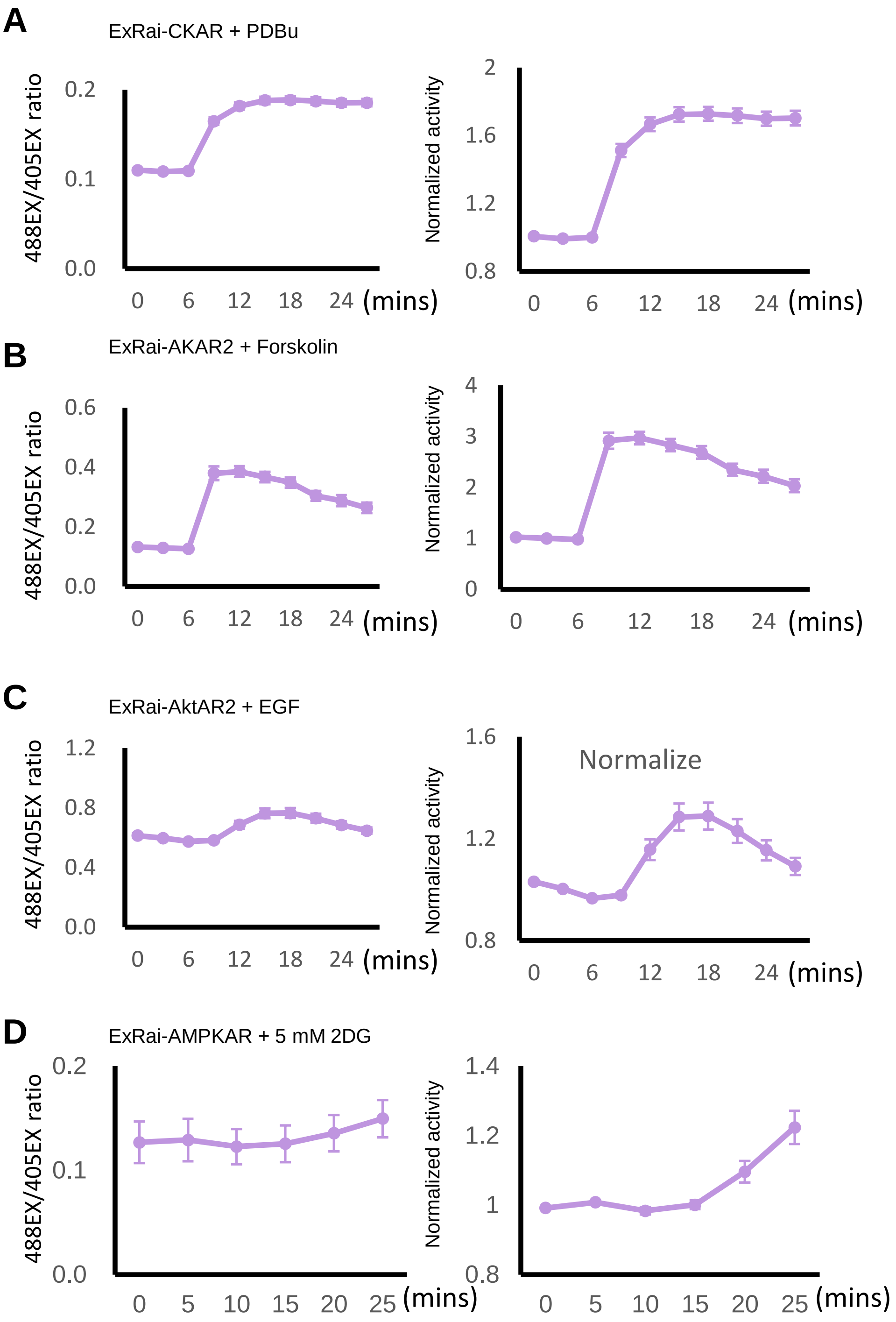


**Figure S4. Responses of ExRai biosensors.** Cells expressing the indicated ExRai biosensors were treated with known activators. The responses (mean+/-SEM of n =20 cells) are normalized to prestimulus levels.
